# Curcumin Inhibition of DYRK Kinases

**DOI:** 10.1101/2024.06.10.598318

**Authors:** Filipe Menezes, Przemyslaw Grygier, Katarzyna Pustelny, Grzegorz M. Popowicz, Anna Czarna

**Author notes:** Correspondence to Anna Czarna. These authors contributed equally (Filipe Menezes, Przemyslaw Grygier).

## Abstract

Curcumin is known as a dietary supplement with several health benefits, including antioxidant, blood sugar-lowering, anti-inflammatory, and anti-cancer properties. These benefits make it useful in the treatment of diabetes, neurodegenerative diseases, and cancer. This research focuses on how curcumin interacts with dual-specificity tyrosine-regulated kinases (DYRKs), particularly DYRK1A and DYRK2, which control important cellular processes such as protein breakdown, gene activity and DNA packaging. DYRK1A is crucial for the growth and survival of pancreatic β-cells, which are key in the treatment of diabetes. Curcumin helps increase insulin release and sensitivity by affecting these cells. DYRK2, on the other hand, is involved in cancer, where curcumin’s ability to block its activity helps to reduce tumour growth and spread. These interactions demonstrate curcumin’s potential as a versatile treatment option. By affecting DYRK1A and DYRK2, curcumin promotes β-cell health in diabetes and fights cancer by inhibiting cell growth and promoting cell death. This study underscores the promise of curcumin as a natural compound for treating complex diseases associated with oxidative stress and inflammation, highlighting the need for a better understanding of DYRK kinases to effectively exploit their role in disease processes and cell signalling.

## Introduction

Curcumin, diferuloylmethane, is a naturally occurring polyphenol and the principal component of turmeric, sourced from the rhizome of *Curcuma longa*. The adoption of phytochemicals like curcumin has revolutionised the pharmaceutical landscape, offering low-cost, widely available treatments with minimal side effects. Over the past two decades, extensive research has revealed curcumin’s diverse biological activities, underscoring its therapeutic potential for a broad range of health conditions.

Curcumin plays a crucial role in traditional and alternative medicines, especially in Ayurveda and traditional Chinese medicines, where it is used to treat many health conditions, ranging from minor ailments to complex diseases [1]. Additionally, curcumin has gained popularity as a dietary supplement and nutraceutical, becoming one of the most successful natural health products sold in the United States since 2013 [2]. Renowned for its antioxidant, hypoglycemic, anti-inflammatory, and anti-cancer properties, curcumin is turning into a versatile therapeutic agent. It modulates a range of signalling molecules, including transcription factors, chemokines, cytokines, tumour suppressor genes, adhesion molecules, and microRNAs. To date, over 100 different clinical trials have been completed with curcumin proving its safety, tolerability, and effectiveness in addressing various chronic diseases in humans [3]. Numerous scientific reports emphasise its antidiabetic effects based on cellular studies, animal models, and clinical trials [4], [5], [6], as well as its potential implications in neurodegenerative disorders and cancer. Unsurprisingly, curcumin is drawing the interest of the scientific community.

Of particular interest is curcumin’s protective effect against Type 2 Diabetes (T2D), which is thought to be mediated through a direct influence of pancreatic β-cells. This promotes insulin secretion and improves glucose regulation. To effectively treat diabetes, therapeutic strategies must restore cellular homeostasis but also target the regeneration of pancreatic β-cells. Potential therapeutic targets for influencing the β-cell cycle include the glucagon-like peptide-1 (GLP-1) receptor [7], DYRK1A [8], transforming growth factor-β receptor (TGF-βR) [7], glycogen synthase kinase-3β (GSK3β) [9], the phosphatidylinositol 3 kinase (PI3K)-serine-threonine protein kinase B (Akt)-mammalian target of rapamycin (mTOR) signalling pathway [10], and serotonin receptor 2B (HTR2B) [11]. Currently, only GLP-1R agonists are available on the market, but their impact on human β-cell proliferation is limited. Further exploration into the dual inhibition of DYRK1A and the regulatory roles of DYRK2 by curcumin and its derivatives could lead to new therapeutic strategies that enhance β-cell regeneration and mitigate diabetes-related complications. Integrating these findings into comprehensive treatment approaches may significantly improve outcomes for individuals with T2D.

Conversely, DYRK2 has been predominantly studied in the context of cancer, where it regulates cell proliferation, apoptosis, and DNA damage responses. Significant studies have highlighted DYRK2’s role in various processes. Recent research has demonstrated that DYRK2 regulates the epithelial-mesenchymal transition (EMT) in breast cancer by promoting the degradation of SNAIL, a key transcription factor in EMT and cancer metastasis [12]. Another study elucidates the structural basis for DYRK2 inhibition by curcumin, revealing the binding mode and its kinase activity inhibition [13]. DYRK2’s role in DNA damage response and its potential as a therapeutic target in cancer treatment has also been highlighted. This emphasises the role of this kinase in maintaining genomic stability [14]. Targeting DYRK2 with small molecules like curcumin holds significant therapeutic potential in cancer and other diseases [15].

## Results and Discussion

### THE STRUCTURE OF CURCUMIN IN DYRK1A

The DYRK1A-curcumin complex was crystallised in the P 63 space group with 2 molecules in the asymmetric unit. The structure was solved at 2.69 Å resolution. The molecules comprising the asymmetric unit superimpose with a root-mean-square deviation (rmsd) of 0.565 Å over 304 C_α_ atoms.

Both chains contain a clearly interpretable electron density of curcumin in the ATP-binding pocket. One of the aromatic rings of the molecule is located close to the kinase’s catalytic centre and π-stacks with Phe238. The methoxy and phenol groups attached to this aromatic ring provide two optimally oriented hydrogen bonds (H-bonds) with Lys188 and Glu203. This hydrogen bonding pattern is similar to the one reported for the DYRK2-curcumin complex (see PDB-ID 6HDR). In addition to taking part in the hydrogen bonding with the aminium group of Lys188, the methoxy group might also contribute to binding by forming lipophilic contacts with the sidechains of Phe170, Val173, and the methylene groups of Lys188’s side chain.

The dione group of curcumin is expected to be in equilibrium with an enol tautomer, which seems to be the predominant species bound to DYRK1A. One of the oxygens from this group forms an H-bond with a main chain carbonyl of the hinge residue Glu239 and another with the main chain nitrogen of Leu241. These interactions resemble a β-strand organisation in proteins. Nonpolar surfaces of the dione group seem to form a hydrophobic contact with the side chains of Ala186 and Leu294. Interestingly, curcumin does not explore interactions with Ser242, which have previously been shown in cocrystal structures of other DYRK1A inhibitors. Instead, the 3-methoxy-4-hydroxyphenyl group proximal to the entrance of the DYRK1A ATP-binding cavity shields Ser242 from the solvent and forms lipophilic interactions with Met240, Ile165, and a PEG molecule adjacent to the binding cleft. The latter results from the cryoprotectant solution used during the cryocooling of the crystals. In the DYRK2 structure, the methoxy group of curcumin forms a hydrogen bond with Lys153. In contrast, in the DYRK1A structure, there is no such interaction since Ser163 occupies the same position. The lack of direct interactions of the 3-methoxy-4-hydroxyphenyl group in DYRK1A allows for additional flexibility. Comparing both molecules in the asymmetric unit reveals opposite orientations: the ring is flipped by 180 degrees with the methoxy moiety pointing in opposite directions. In contrast to most of the published DYRK1A inhibitors, curcumin appears to bind without bridging water molecules. Moreover, it does not utilise the aromatic rings to reach the depth of the binding cleft, as seen, for example, in the structure with harmine or flavone inhibitors.

As already mentioned, the 3-methoxy-4-hydroxyphenyl group in the proximity of the catalytic centre of the kinase forms interactions similar to those described for the DYRK2-curcumin complex. However, the dione moiety appears to be preferentially bound in DYRK1A as the distances between the H-bonding atoms of the protein and ligand are much closer (2.6 and 2.64 A to Glu239 and Leu241, respectively) compared to DYRK2 (3.29 and 2.92 to Glu229 and Leu231, respectively). Moreover, Met240 seems to have more extensive lipophilic interactions than the Leu230 of DYRK2. Therefore, the interaction with DYRK1A is more specific as it relies primarily on polar group complementarity.

### QUANTUM MECHANICAL COMPARISON OF CURCUMIN-DYRK COMPLEXES

Although curcumin shows low affinity towards the DYRK kinases, data shows a preference towards the paralog DYRK2 [16]. When compared to DYRK1A, a difference of at least an order of magnitude is observed in the IC_50_s. This prompted us to further investigate the underlying molecular mechanisms that could help rationalise and improve selectivity and activity using our Energy Decomposition and Deconvolutional Analysis (EDDA) [17], [18]. Figure 2 schematises the binding mode of curcumin to DYRK2 and DYRK1A and shows the most relevant interaction maps. Figure 3 represents the differential EDDA plot, quantifying differences in the nature of protein-curcumin interactions.

**Figure 1:**
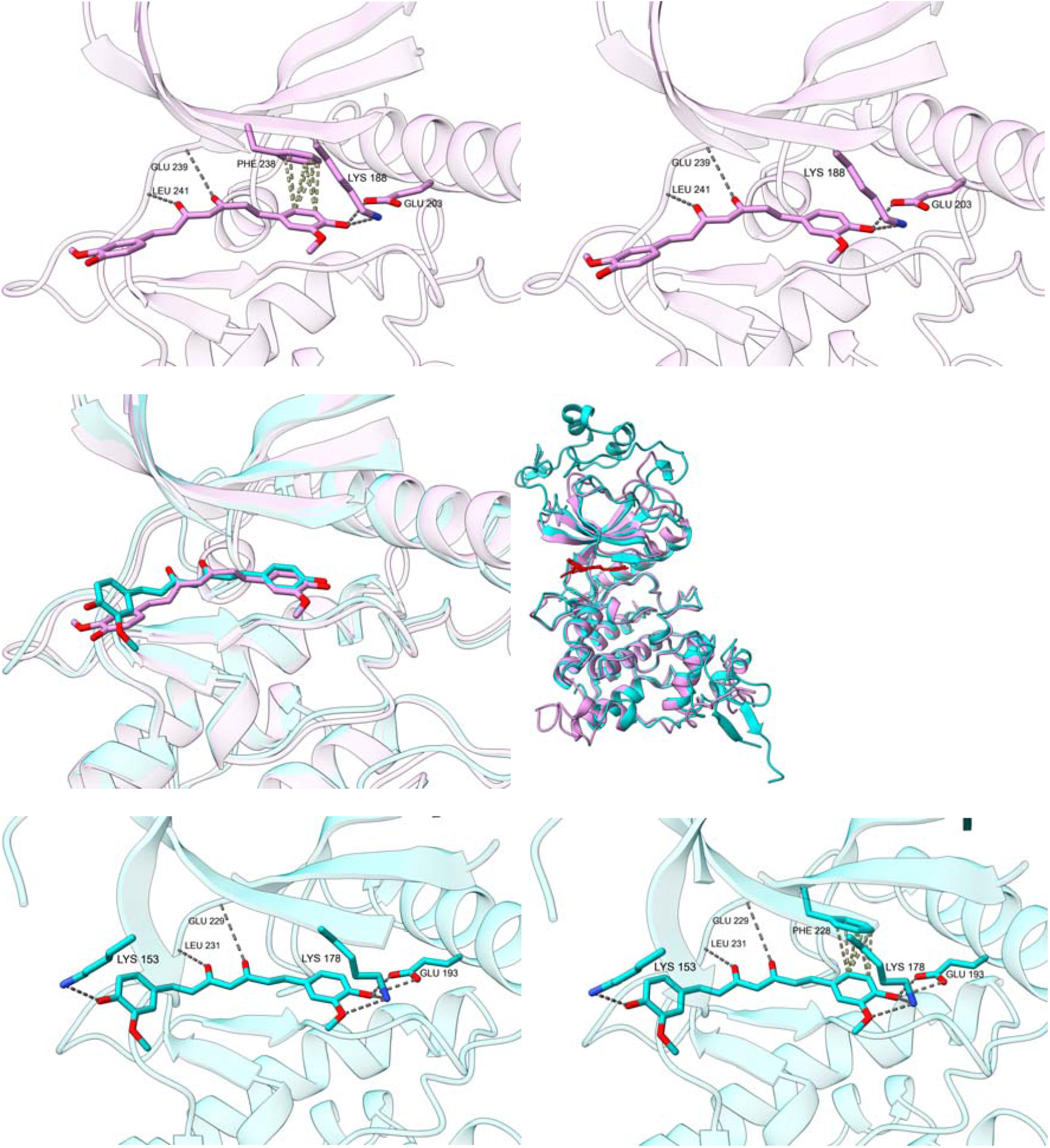
The binding pose of curcumin in DYRK1A and DYRK2.

**Figure 2.**
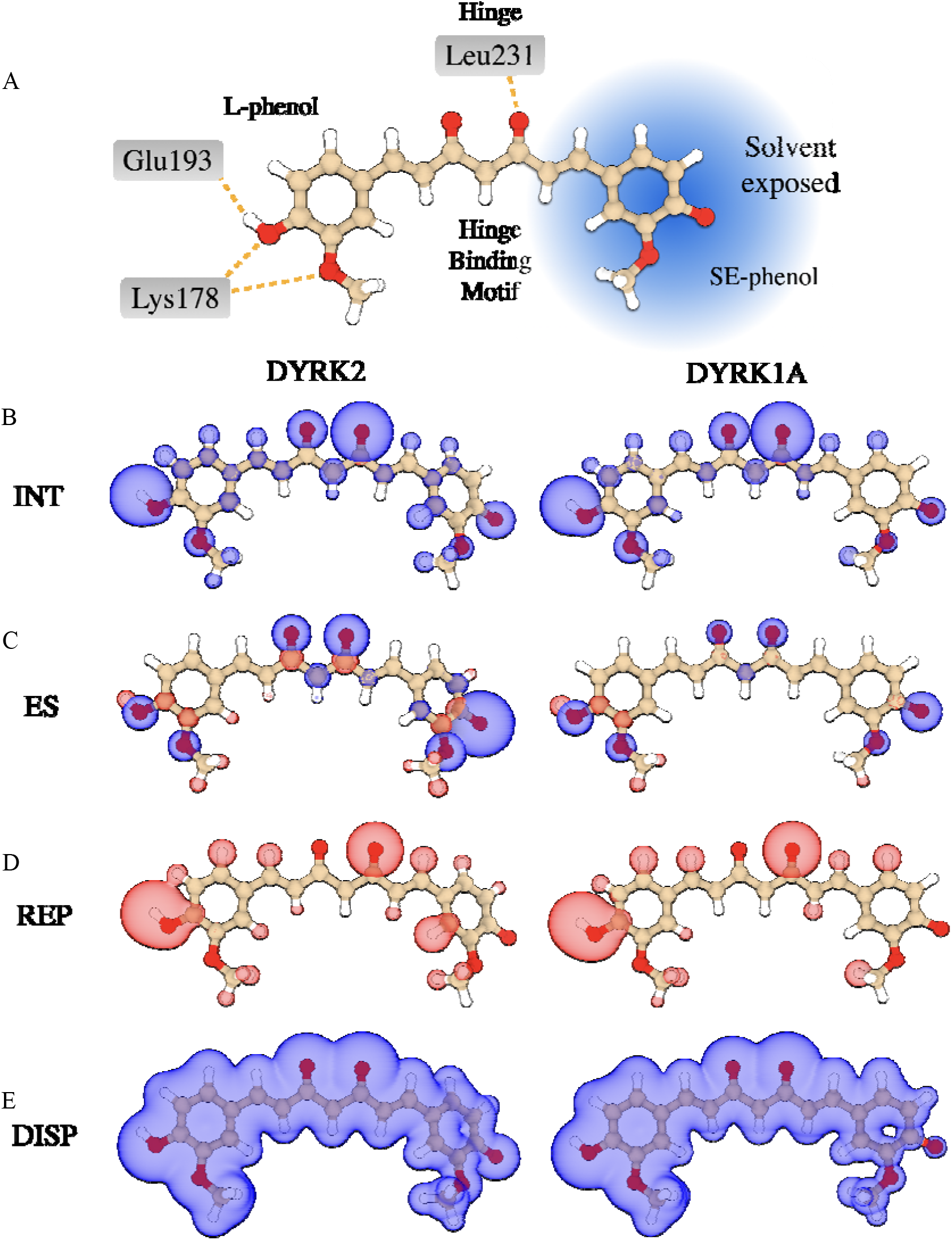
Comparison for curcumin binding in DYRK2 and DYRK1A. **A)** Schematic representation of how curcumin binds each of the kinases. Residue numbering according to the DYRK2 structure (6HDR). The interaction pattern with the pocket is identical for DYRK1A. However, the residue numbering is shifted by 10 (*e*.*g*., instead of Leu231, the interaction takes place with Leu241). **B)** EDDA *imaps* for the binding energy of curcumin in DYRK2 (left) and DYRK1A (right). *Imaps* for **C)** electrostatic interactions, **D)** electronic density repulsion, and **E)** dispersion forces or lipophilicity.

**Figure 3.**
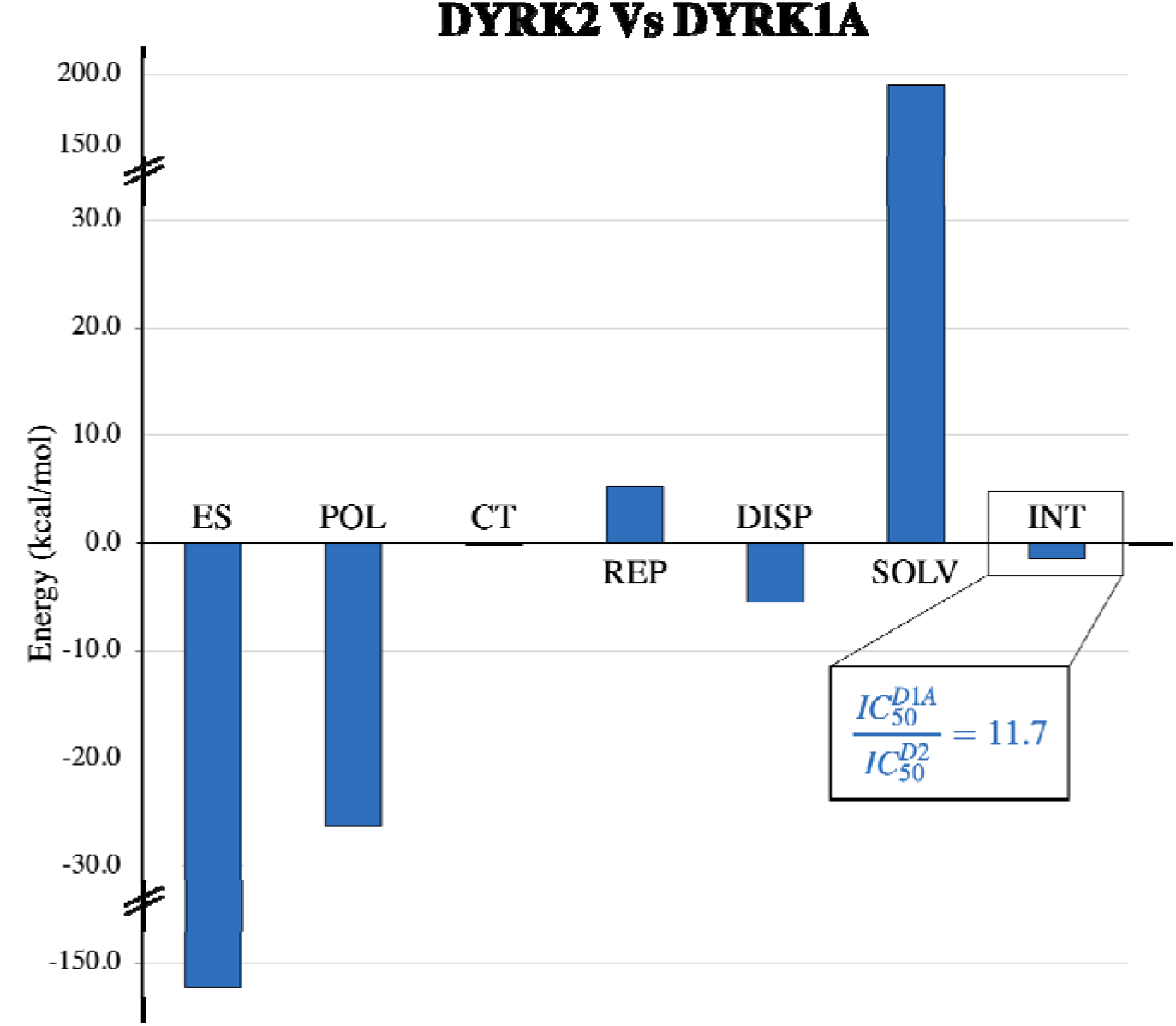
Differential EDDA plot for curcumin in DYRK2 and in DYRK1A. All bar plots taking negative values mean that interaction type is favoured for DYRK2. From the difference in binding energies, we estimate a relative IC_50_ of 11.7, favouring DYRK2. This agrees with experimental data available in the literature [16]. ES stands for electrostatic interactions, POL for exchange-polarization, CT is charge transfer, REP is the repulsion of electronic clouds, DISP dispersion forces, SOLV the solvation contributions, and INT the binding energy.

The calculations indicate a lower binding energy for DYRK2 than for DYRK1A (**Figure 3**). The difference in energy is such that the relative dissociation constants are about a factor of 11.7 larger for DYRK1A, in excellent agreement with published data [16]. The total interaction maps reveal a strong, attractive contribution by the solvent-exposed phenol group (SE-phenol) and another one by the other phenol, binding to the catalytic Lys and Glu193 (Glu203 in DYRK1A; L-phenol; **Figure 2B**). The hinge binding group, characterised by two carbonyls or the respective enolate, shows an equally strong interaction in both cases. Electrostatics, polarisation, and dispersion (lipophilic) interactions are overall favouring the DYRK2 paralog. The maps for electrostatic interactions, **Figure 2C**, reveal a particularly stabilising interaction involving the SE-phenoxy. This can be traced back to an H-bond and an ionic bridge with the nearby Lys153. Note that this interaction is absent in the case of DYRK1A, which exposes a leucine in position 163 (equivalent to 153 in DYRK2). However, in both kinases, there is a water-mediated interaction with the more distal Lys175. The lipophilicity maps, **Figure 2E**, further reflect the close contact explored by the water molecule, Lys153, and the SE-phenoxy group of curcumin when binding to DYRK2. Similar observations may be taken from the total interaction maps.

Electrostatics also evidence differences in how the carbonyls bind to the hinge (Leu231 in DYRK2, Leu241 in DYRK1A), again favouring interactions for DYRK2. The repulsion maps shown in **Figure 2D** indicate clearly that the interaction mode is conserved between curcumin and the two kinases. When analysing the total interaction maps, there is a slight preference towards DYRK1A in the hinge binding motif, which indicates that an additional factor is cancelling the electrostatic stabilisation brought by the two carbonyls. This may be found in the desolvation penalties required when binding occurs (see supplemental, Figure SC1). The calculations indicate a stronger electrostatic character for DYRK2. This can be easily understood given the higher density of charged residues in the protein.

The calculations also indicate a slight preference for curcumin’s L-phenol group to bind the nearby Glu residue of DYRK2 (see **Figure 2A**). This is reflected in the fine details of the repulsion maps. Electrostatic interactions between the inhibitor and the kinases are, in this case, visually indistinguishable. The charge transfer and polarisation maps indicate stronger contributions in the case of the interaction between curcumin and DYRK2 (see supplemental Figure SC1). This seems to indicate a slightly stronger covalent character in this hydrogen bond. However, the effect is expected to be minimal since differences in charge transfer contributions are negligible. Note, however, that the interaction lobe is distorted towards the oxygen atom in the case of DYRK1A, indicating a stronger contact with the catalytic lysin rather than the nearby Glu residue. Lastly, the calculations show a slightly stronger attractive interaction surface between curcumin and DYRK2. This is seen in the larger number of attractive contacts between inhibitor and target, extending, in the case of DYRK2, to the whole π -system.

When analysing the computational data, a clear statement seems to emerge, in which the selectivity of curcumin to DYRK2 should result from a more efficient interaction surface that promotes the lipophilic contacts with curcumin’s π -system and capturing an H-bond with Lys153. In particular, the latter seems to be a key element in ascertaining selectivity between the two kinases. To further corroborate our hypothesis, we compared the sequences of DYRK1B [19] and DYRK3 (PDB 5Y86) [20] to those of DYRK1A and DYRK2. IC_50_ data on DYRK1B shows that this kinase is as poorly inhibited by curcumin, much like the case of DYRK1A [16]. Common to both kinases is the absence of lysin at the equivalent of position 153 in DYRK2. On the other hand, the IC_50_ of curcumin in DYRK3 is *ca*. 4 times larger than that of DYRK2. At the equivalent of position 153, DYRK3 also places a lysine. Note that a relative IC_50_ of 4 is equivalent to a relative difference in Gibbs free energy of *ca*. 0.8 kcal/mol, which could easily be accounted for by fine details of the π interaction surface.

### QUANTUM MECHANICAL ANALYSIS OF CURCUMIN-DYRK1A BINDING MODE

The crystal of curcumin in DYRK1A contains two protein-ligand complexes in the asymmetric unit. These show the inhibitor in different binding conformations: in one, the ligand is linearly placed, constituting the pose we compared against the structure published for curcumin in DYRK2; in the additional complex, curcumin bends the solvent-exposed groups (see **Figure 4**). This indicates a certain degree of flexibility of curcumin in the kinase’s pocket, allowing us to retrieve additional cues on the nature of interactions. The relative binding energies show a preference towards the linear binding mode of curcumin (**Figure 5**). This is supported by an attractive contribution to determining the nature of protein-ligand interactions: electrostatics. One may follow in the electrostatics maps (**Figure 4B**) that two of the three main interaction points offer stronger contacts: the hinge binding motif and the phenol targeting the catalytic lysine. The solvent-exposed phenolate seems favoured in the bent pose of curcumin because of a water-mediated interaction with Lys175. However, a few discrepancies arise when we compare the relative strength and the directionality of electrostatic interactions with the total interaction maps (**Figure 4A**). Specifically, the total interaction on the hinge binding motif seems to favour bent curcumin in the kinase’s pocket. This is further supported by the electronic repulsion maps, which evidence a shorter contact to the main chain’s amide proton of Leu241. Again, the fine differences in determining the preferred binding pose seem to lie in the weak complementarity between inhibitor and pocket. This observation is particularly interesting, as it indicates that curcumin lacks any strong or directed interaction site with the kinase, unlike other inhibitors of these kinases. This forms the basis of curcumin’s poor inhibitory power, corroborating that the scaffold lacks any specific feature characteristic of an interesting lead candidate.

**Figure 4.**
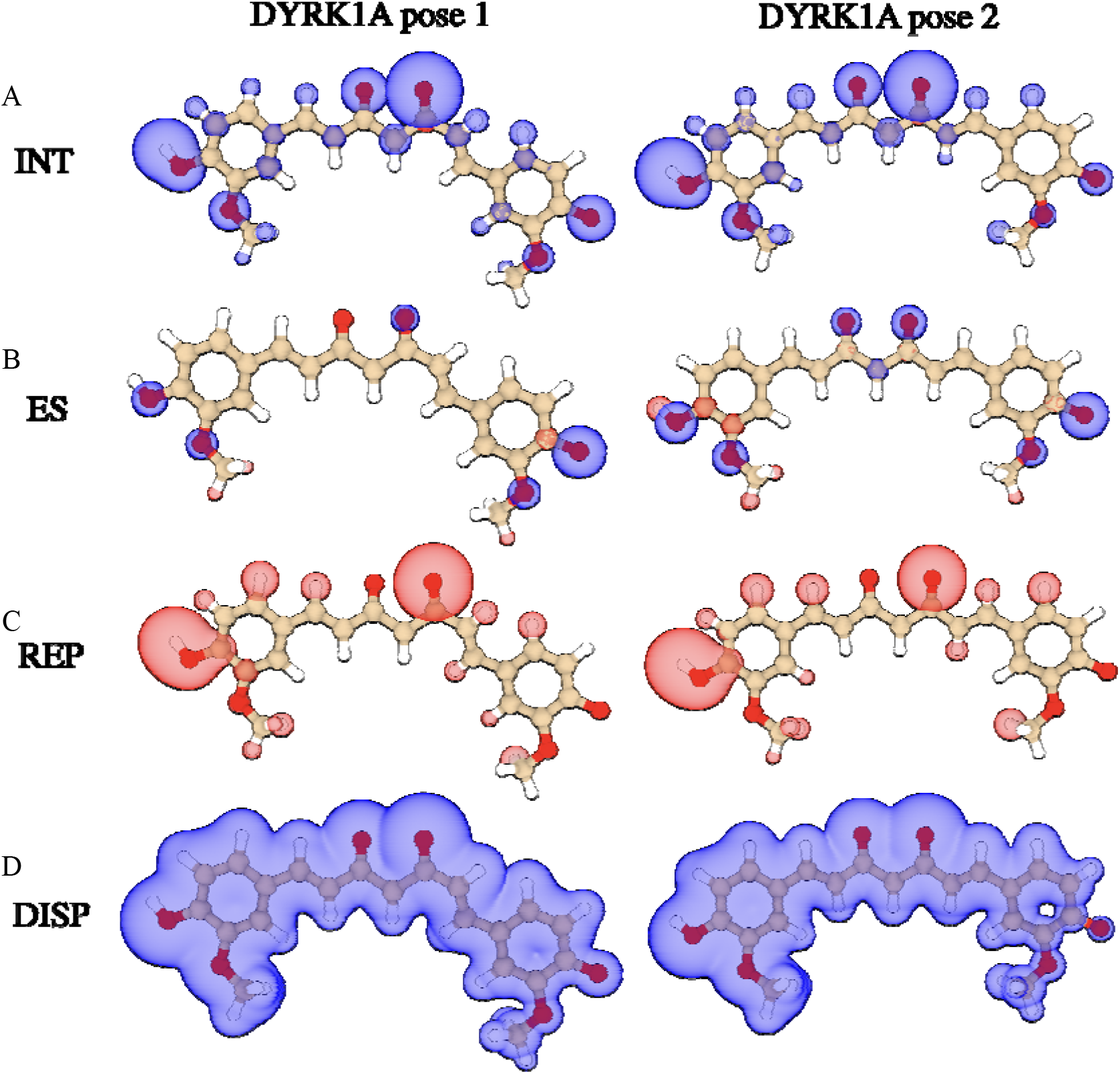
Binding mode comparison for curcumin in DYRK1A. **A)** EDDA imaps for the binding energy of curcumin’s bent (left) and linear (right) poses in DYRK1A. Imaps for **B)** electrostatic interactions, **C)** electronic density repulsion, and **D)** dispersion forces or lipophilicity.

**Figure 5.**
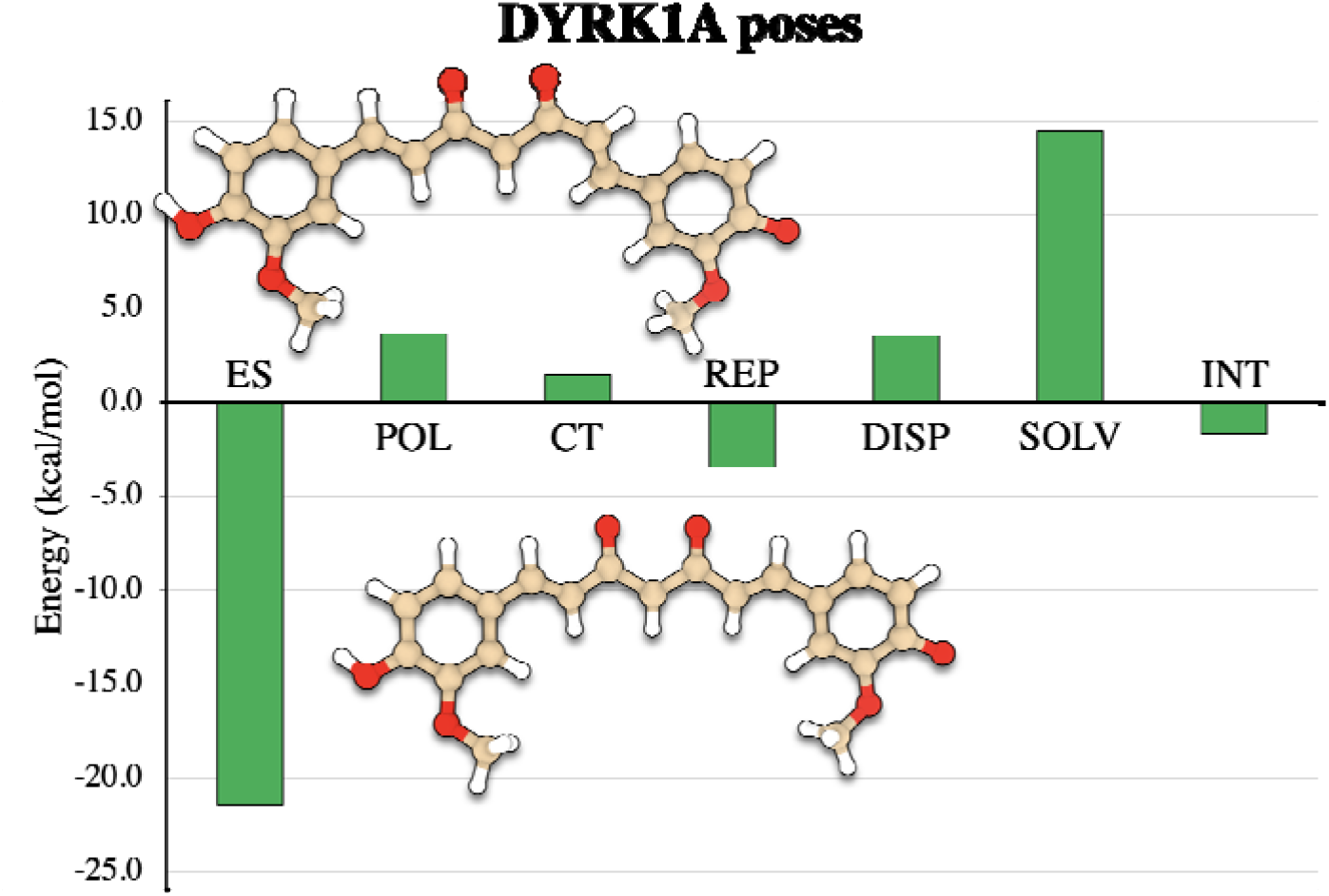
Differential EDDA plot for curcumin’s binding poses in DYRK1A. All bar plots taking negative values mean the interaction type favours the linear pose. The calculations indicate an overall preference for the linear pose.

Besides the observed conformational freedom, curcumin offers the possibility to bind the kinases in different protonation states. Specifically, the solvent-exposed phenolate may capture a proton (the estimated pKa for the free molecule is 9.54 [21]). Also, the enolate binding to the hinge may be protonated (pKa = 8.79). Note that the phenol group binding to the catalytic lysin will likely be protonated; otherwise, a strong repulsion will arise with Glu193/203. Note also that the previously presented data in this manuscript assumed curcumin in a doubly deprotonated state.

Calculations using different protonation states (**Figure 6** and Supplementary tables) systematically show an increase in binding energy upon protonation of the inhibitor (less favourable). On the hinge binding motif, the rationale is clear. By protonating the enolate, its ability to H-bond Leu231/241’s main chain amide is significantly hampered. Questions could potentially arise for the solvent-exposed phenol group because this is reasonably far from the centre of the ATP binding pocket, but also due to its larger pK_a_. However, the calculations indicate a significant gain in binding energy due to electrostatics, but the relative binding energies also come closer in agreement with experimental affinity data. Interestingly, deprotonation of the solvent-exposed phenol seems less important in the case of DYRK2, as the binding energies for the singly and doubly deprotonated states of curcumin are numerically very close (kcal/mol, favouring the doubly deprotonated structure). This suggests a more complex binding model of curcumin in DYRK2, which brings additional entropy stabilisations because the inhibitor is described by different protonation states when bound to the protein. In the case of DYRK1A, the effect is not expected to be as pronounced.

**Figure 6.**
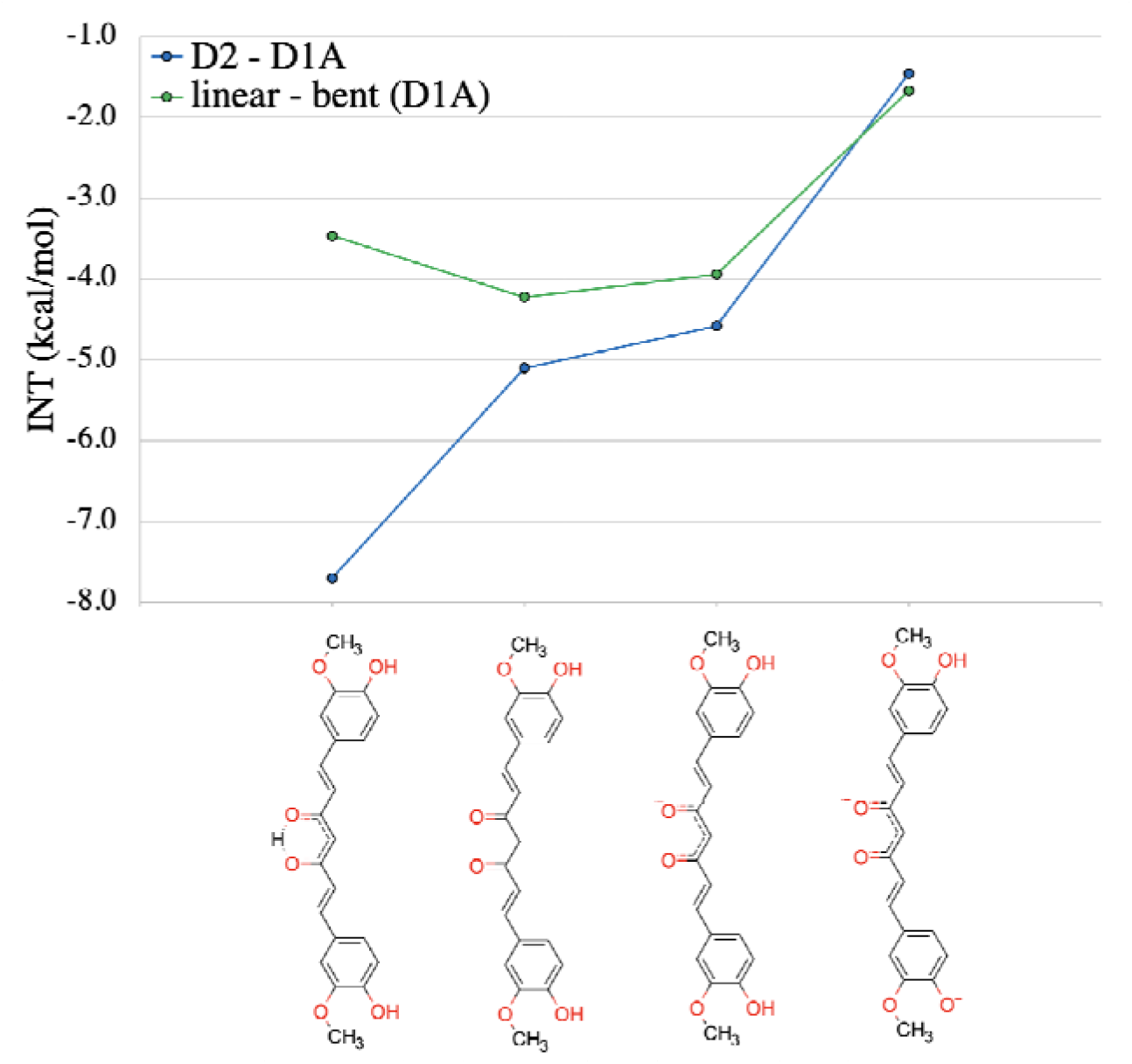
Differential interaction energy for curcumin in DYRK2-DYRK1A (blue) and curcumin’s binding poses in DYRK1A (green) as a function of curcumin’s protonation state.3. The comparison between kinases is assembled in the same fashion as in **Figure 3**, meaning that a negative value shows a preference towards DYRK2. The comparison of the two poses of curcumin in DYRK1A is obtained as in **Figure 5**, meaning. This means that negative values show a preference for the linear binding pose of curcumin. As the total charge in curcumin increases, differences in the binding energy decrease.

## Materials and Methods

Quantum mechanical calculations were performed using the ULYSSES package [22]. The method of choice was GFN2-xTB [23] using ALPB implicit solvation [24]. The solvent of choice is water. Protein structures were processed using MAESTRO [25]. Pockets were manually cut using previously established templates [17].

## Conclusions

The present study provides a structural model of curcumin in DYRK1A, which is analysed in light of a previusly published structure of curcumin with DYRK2. This allows studying how this natural product inhibits the two homologous kinases. Our quantum mechanical analysis gives a deeper insight into the key elements providing selectivity of curcumin to DYRK2. This is an additional electrostatic interaction with Lys153 of DYRK2, which is absent in DYRK1A. This is put in the context of other DYRK kinases, allowing us to place the relative affinities for several of these kinases. The binding mode of curcumin to DYRK1A was also analysed in detail, showing a preference for linearisation of the inhibitor when bound. The calculations further evidence a complex acid-base behaviour for bound curcumin.

