## Supplementary Material for "Curcumin Inhibition of DYRK Kinases"

**Curcumin Inhibition of DYRK Kinases Supplementary Information**

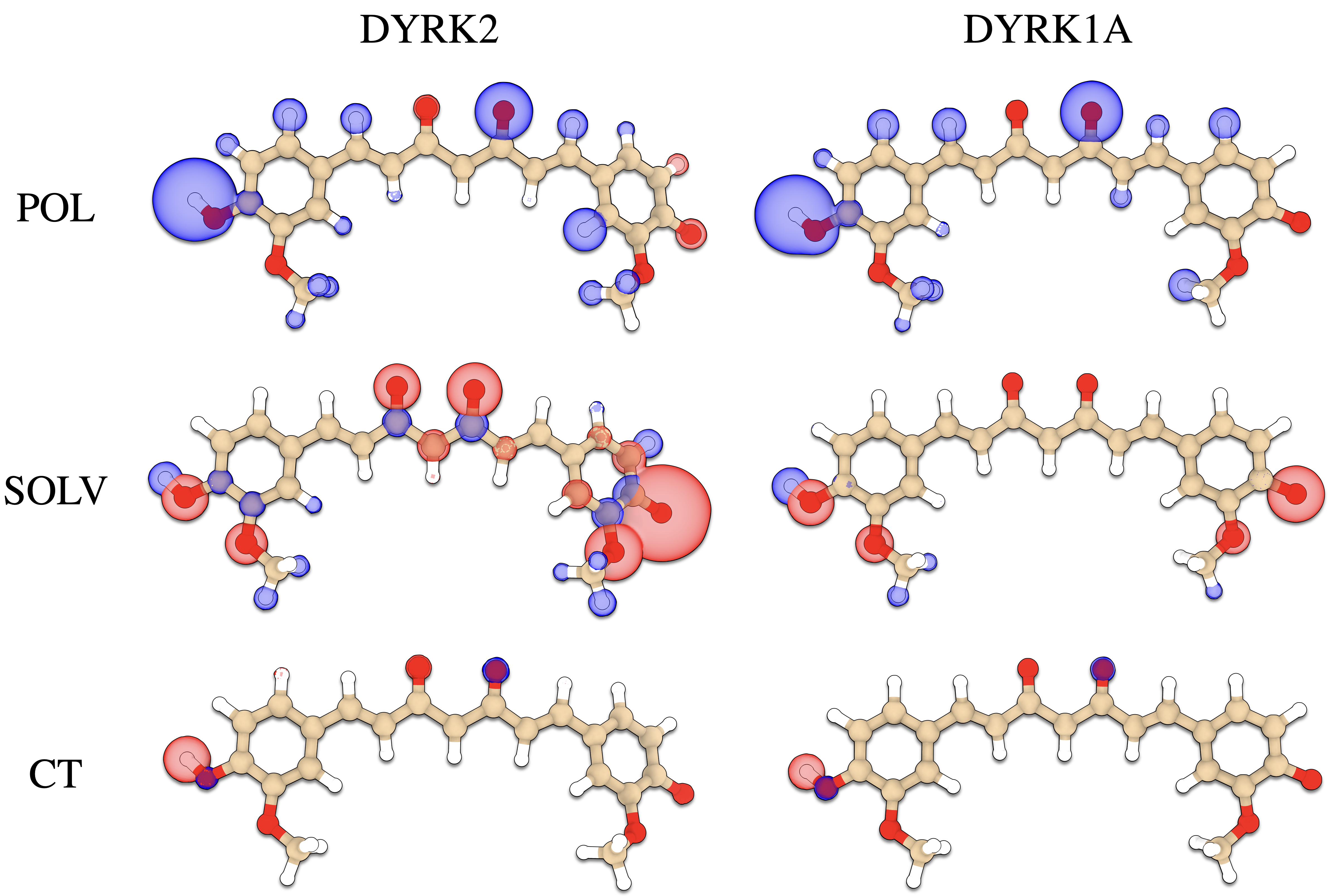

**Figure SC1. Additional *imaps* not presented in Figure C1 for the comparison of curcumin in DYRK2 and DYRK1A.** POL stands for the polarisation energy, SOLV for solvent contribution, and CT for charge transfer.

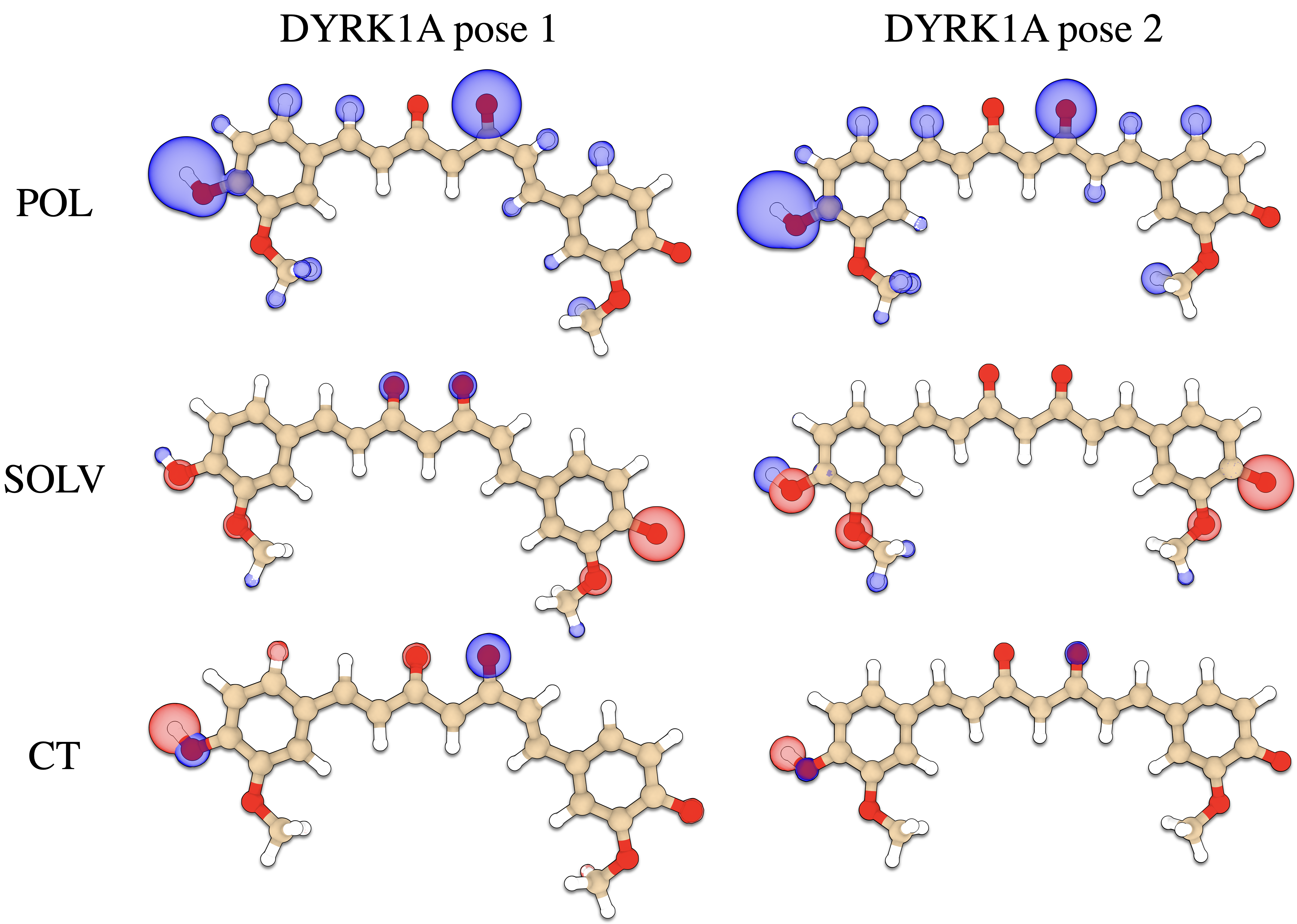

**Figure SC2. Additional *imaps* not presented in Figure C3 for the comparison of curcumin in DYRK2 and DYRK1A.** POL stands for polarization energy, SOLV for solvent contribution, and CT for charge transfer.

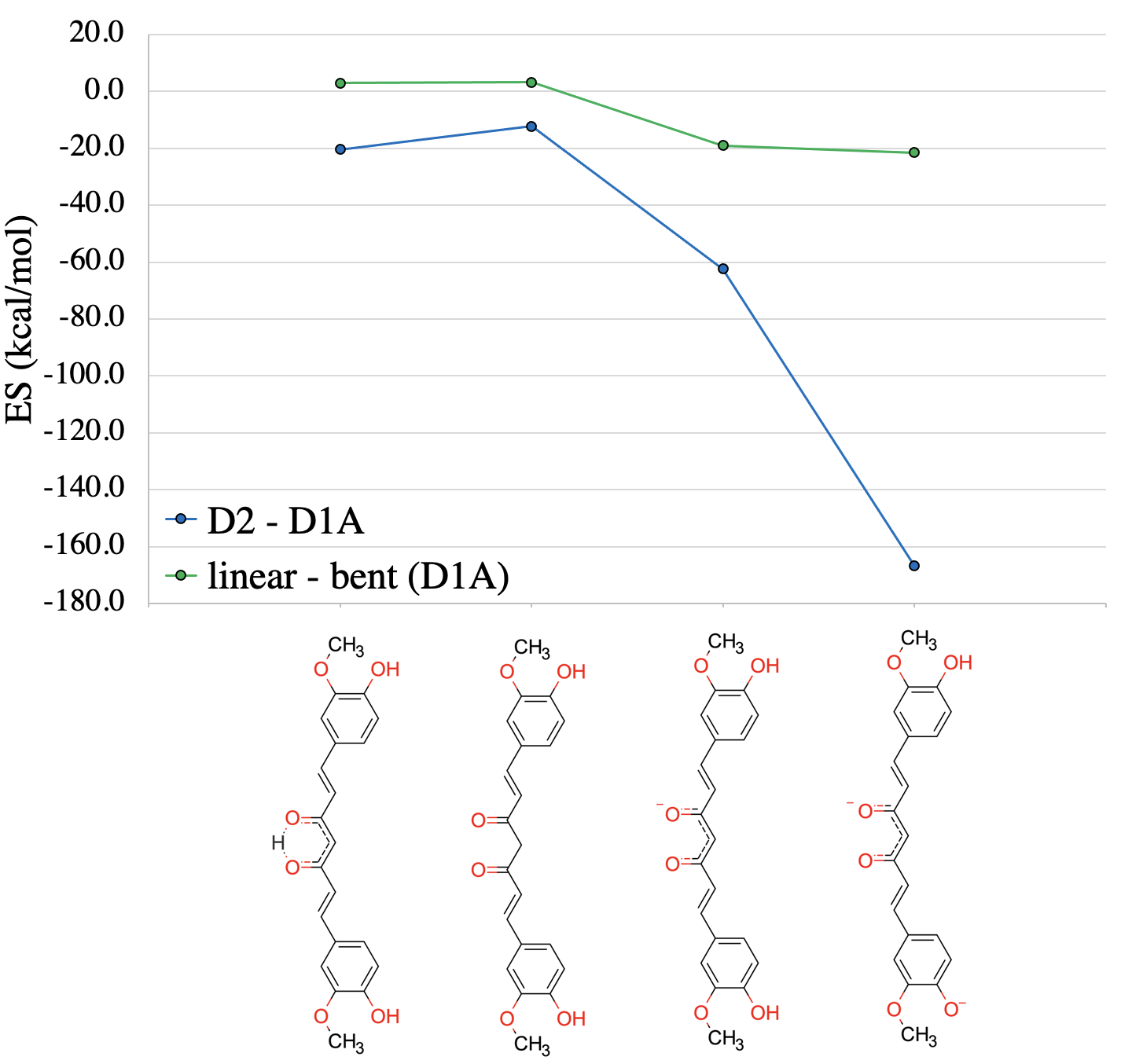

**Figure SC3. Differential electrostatic interaction energy for curcumin in DYRK2-DYRK1A (blue) and curcumin’s binding poses in DYRK1A (green) as a function of curcumin’s protonation state.** The comparison between kinases is assembled in the same fashion as in Figure C2, meaning that a negative value shows a preference towards DYRK2. The comparison of the two poses of curcumin in DYRK1A is obtained as in Figure C4, meaning $\Delta E=linear-bent$. This means that the negative values systematically show a preference for the linear binding pose of curcumin. As the total charge in curcumin increases, differences in the binding energy decrease.

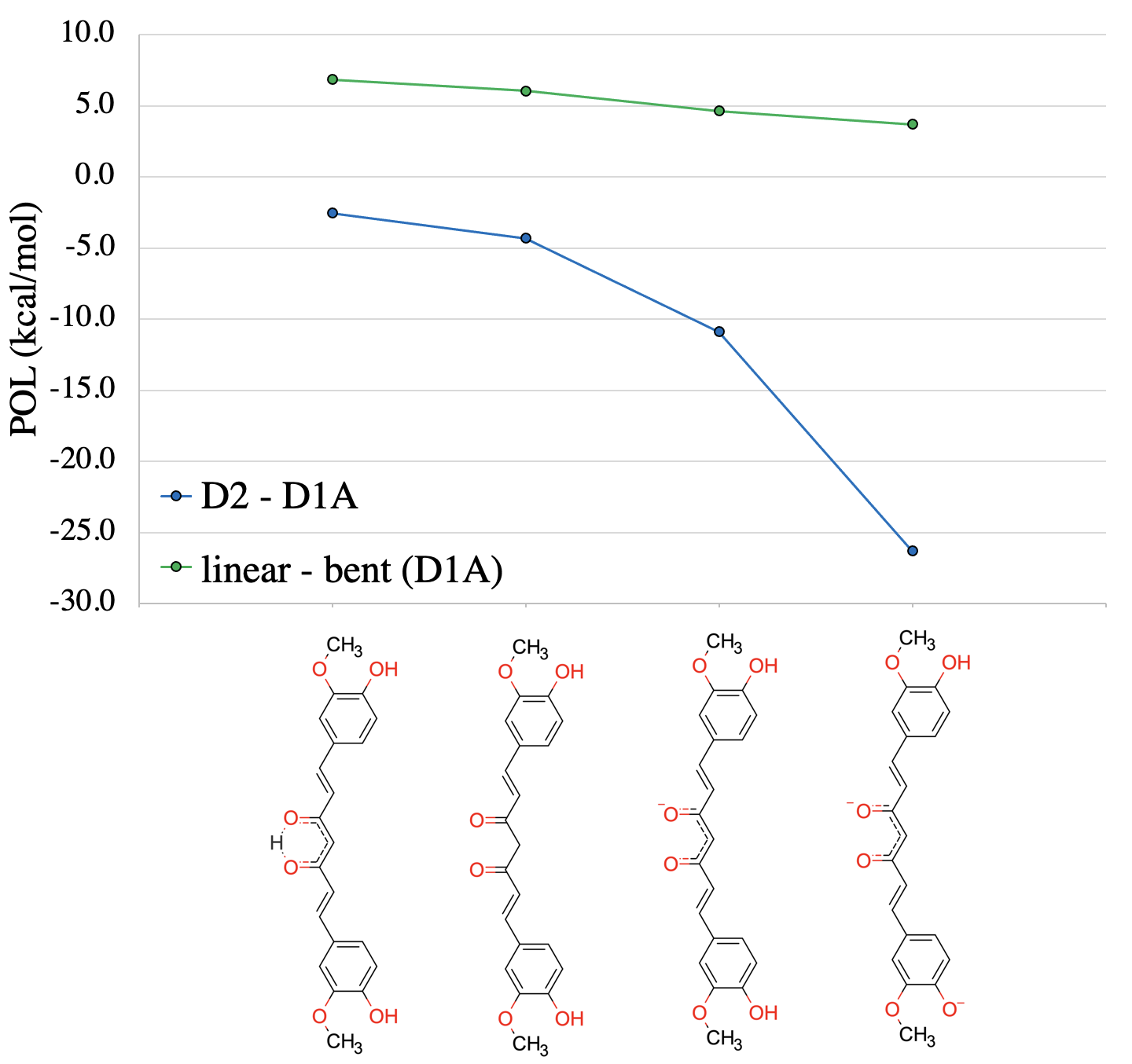

**Figure SC4. Differential polarisation interaction energy for curcumin in DYRK2-DYRK1A (blue) and curcumin’s binding poses in DYRK1A (green) as a function of curcumin’s protonation state.** The comparison between kinases is assembled in the same fashion as in Figure C2, meaning that a negative value shows a preference towards DYRK2. The comparison of the two poses of curcumin in DYRK1A is obtained as in Figure C4, meaning $\Delta E=linear-bent$. This means that the negative values systematically show a preference for the linear binding pose of curcumin. As the total charge in curcumin increases, differences in the binding energy decrease.

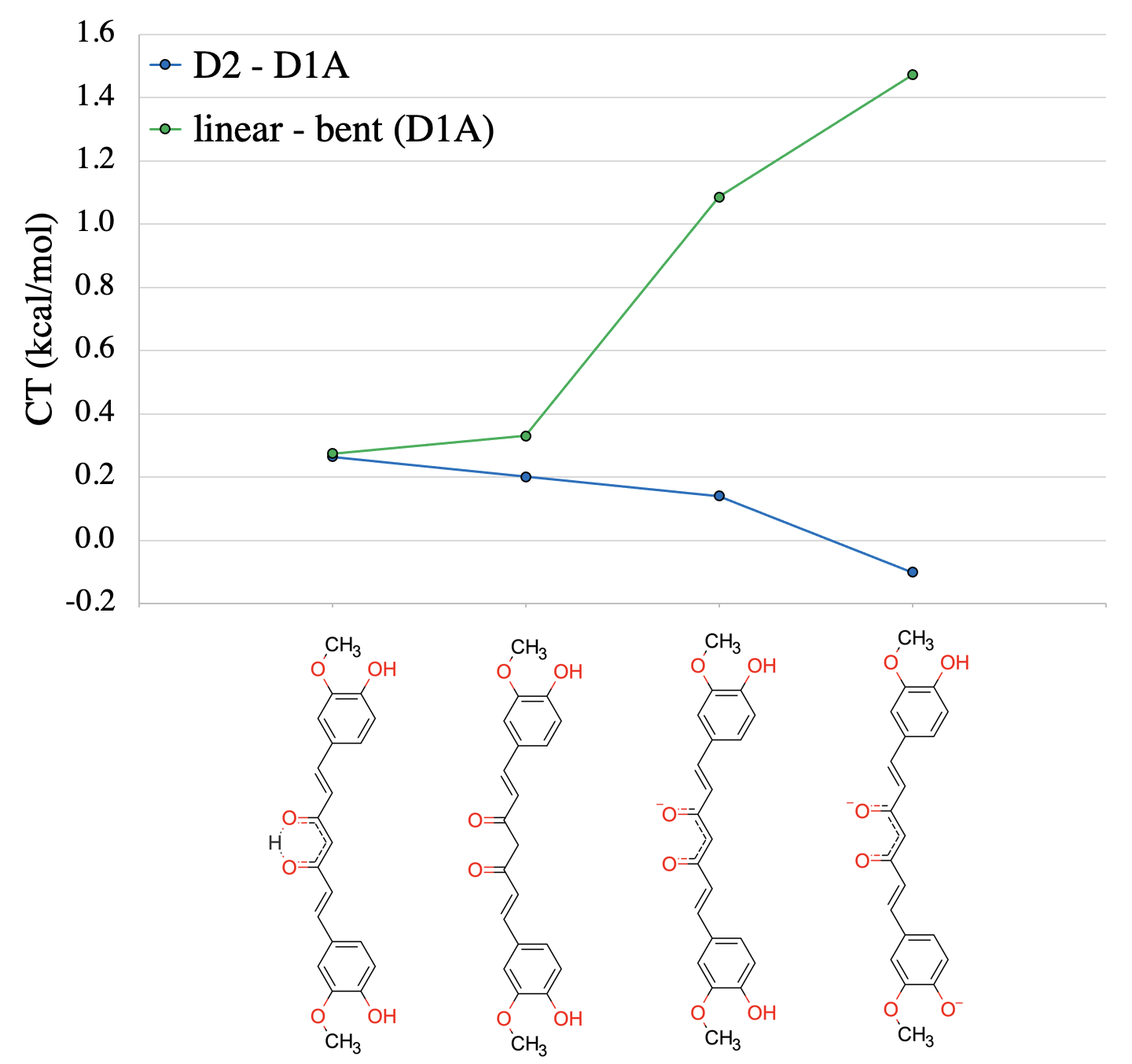

**Figure SC5. Differential charge transfer interaction energy for curcumin in DYRK2-DYRK1A (blue) and curcumin’s binding poses in DYRK1A (green) as a function of curcumin’s protonation state.** The comparison between kinases is assembled in the same fashion as in Figure C2, meaning that a negative value shows a preference towards DYRK2. The comparison of the two poses of curcumin in DYRK1A is obtained as in Figure C4, meaning $\Delta E=linear-bent$. This means that the negative values systematically show a preference for the linear binding pose of curcumin. As the total charge in curcumin increases, differences in the binding energy decrease.

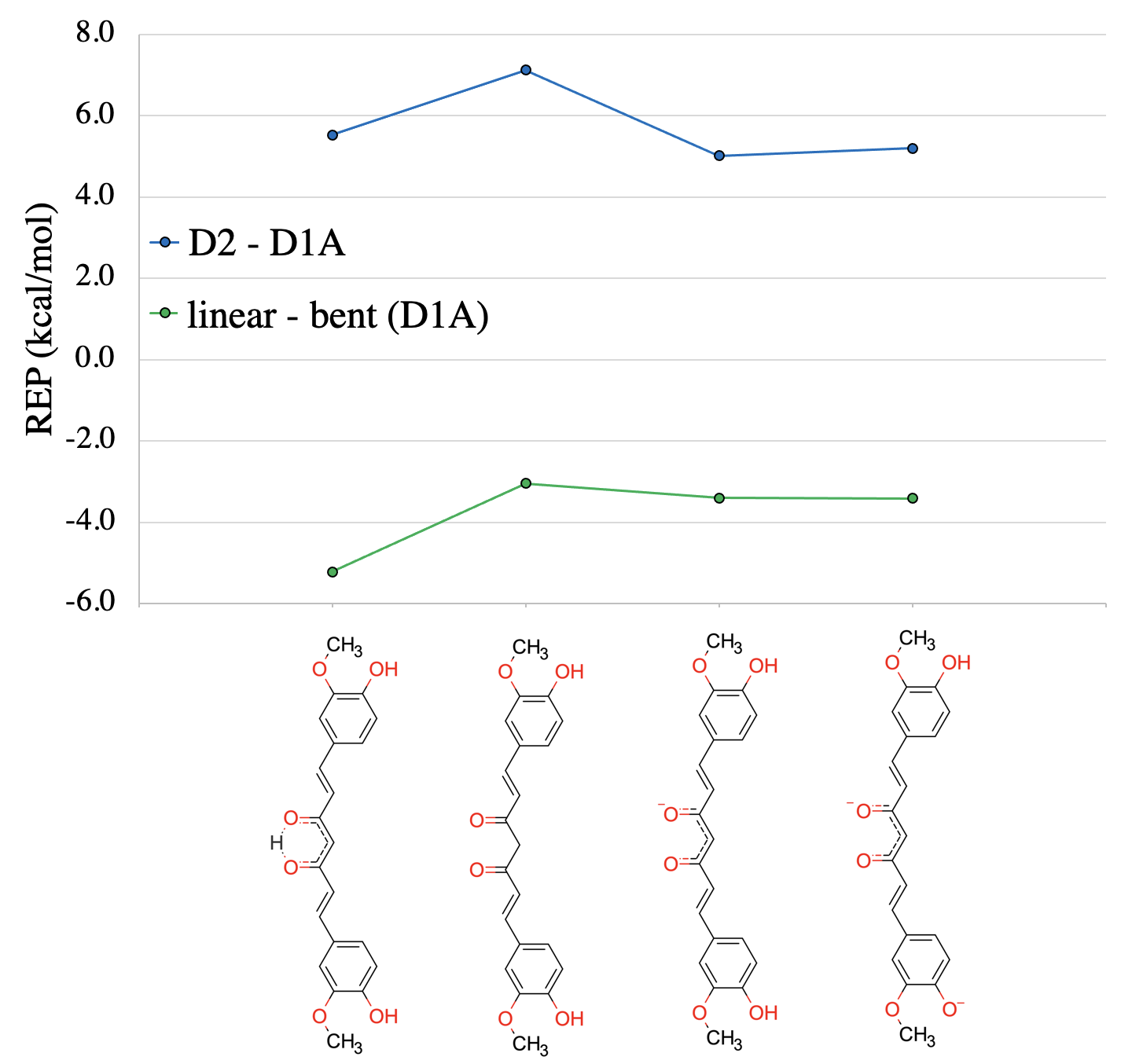

**Figure SC6. Differential electronic density repulsion interaction energy for curcumin in DYRK2-DYRK1A (blue) and curcumin’s binding poses in DYRK1A (green) as a function of curcumin’s protonation state.** The comparison between kinases is assembled in the same fashion as in Figure C2, meaning that a negative value shows a preference towards DYRK2. The comparison of the two poses of curcumin in DYRK1A is obtained as in Figure C4, meaning $\Delta E=linear-bent$. This means that the negative values systematically show a preference for the linear binding pose of curcumin. As the total charge in curcumin increases, differences in the binding energy decrease.

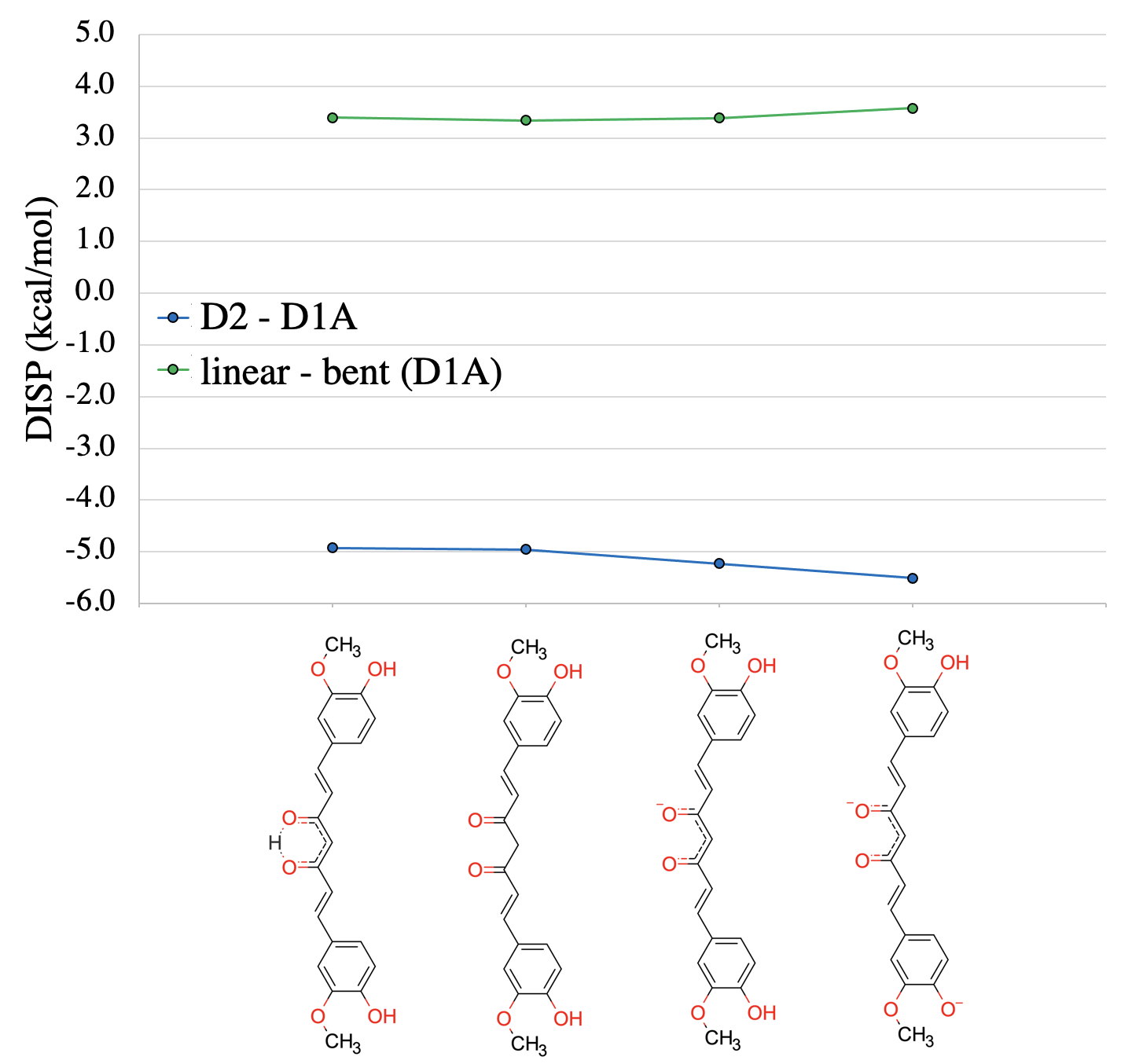

**Figure SC7. Differential dispersion interaction energy for curcumin in DYRK2-DYRK1A (blue) and curcumin’s binding poses in DYRK1A (green) as a function of curcumin’s protonation state.** The comparison between kinases is assembled in the same fashion as in Figure C2, meaning that a negative value shows a preference towards DYRK2. The comparison of the two poses of curcumin in DYRK1A is obtained as in Figure C4, meaning $\Delta E=linear-bent$. This means that the negative values systematically show a preference for the linear binding pose of curcumin. As the total charge in curcumin increases, differences in the binding energy decrease.

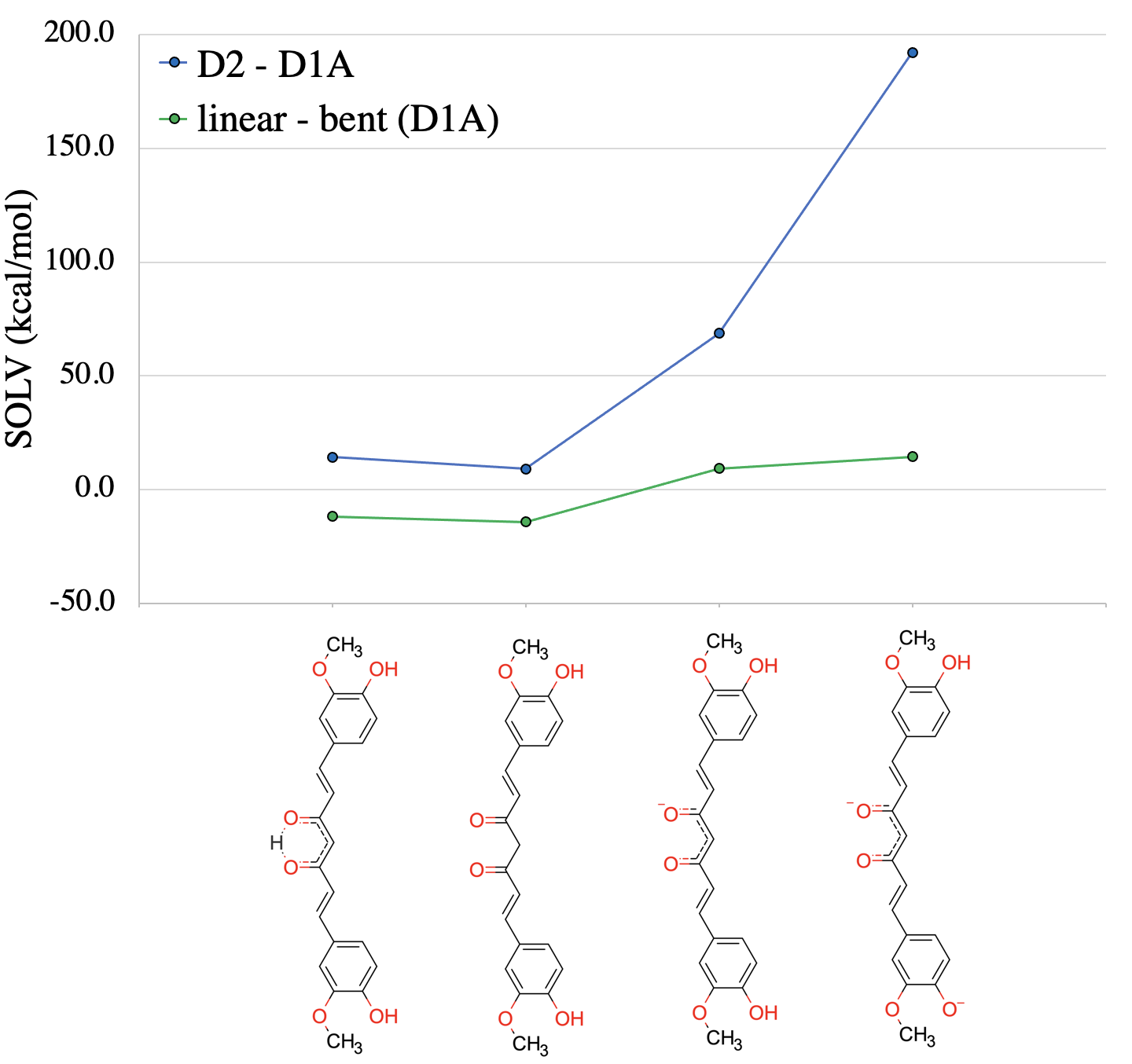

**Figure SC8. Differential solvation energy for curcumin in DYRK2-DYRK1A (blue) and curcumin’s binding poses in DYRK1A (green) as a function of curcumin’s protonation state.** The comparison between kinases is assembled in the same fashion as in Figure C2, meaning that a negative value shows preference towards DYRK2. The comparison of the two poses of curcumin in DYRK1A is obtained as in Figure C4, meaning $\Delta E=linear-bent$. This means that the negative values systematically show a preference for the linear binding pose of curcumin. As the total charge in curcumin increases, differences in the binding energy decrease.

**Table C1. Decomposed energies for curcumin in DYRK2.** The header of the table is encoded according to the total charge (0, 1 – where the enolate is formed, 2 – where the solvent-exposed phenol group is deprotonated). The additional letter code *e* indicates the enol form of the hinge binding motif, whereas *k* marks the respective keto form.

|  | 0*e* | 0*k* | 1 | 2 |
| --- | --- | --- | --- | --- |
| ES | -48.7 | -31.8 | -129.1 | -292.3 |
| POL | -85.8 | -81.3 | -49.6 | -42.1 |
| CT | 0.3 | 0.2 | -0.3 | -0.8 |
| REP | 92.1 | 94.1 | 91.3 | 91.7 |
| DISP | -44.0 | -44.3 | -47.5 | -49.0 |
| SOLV | 37.9 | 16.4 | 84.1 | 241.0 |
| INT | -48.2 | -46.6 | -51.2 | -51.5 |

**Table C2. Decomposed energies for curcumin in DYRK1A, binding pose 2.** The header of the table is encoded according to the total charge (0, 1 – where the enolate is formed, 2 – where the solvent-exposed phenol group is deprotonated). The additional letter code *e* indicates the enol form of the hinge binding motif, whereas *k* marks the respective keto form.

|  | 0*e* | 0*k* | 1 | 2 |
| --- | --- | --- | --- | --- |
| ES | -28.3 | -19.6 | -66.7 | -125.4 |
| POL | -83.2 | -77.0 | -38.7 | -15.8 |
| CT | 0.0 | 0.0 | -0.5 | -0.7 |
| REP | 86.5 | 87.0 | 86.3 | 86.5 |
| DISP | -39.1 | -39.3 | -42.3 | -43.5 |
| SOLV | 23.6 | 7.3 | 15.3 | 48.9 |
| INT | -40.5 | -41.5 | -46.7 | -50.0 |

**Table C3. Decomposed energies for curcumin in DYRK1A, binding pose 1.** The header of the table is encoded according to the total charge (0, 1 – where the enolate is formed, 2 – where the solvent-exposed phenol group is deprotonated). The additional letter code *e* indicates the enol form of the hinge binding motif, whereas *k* marks the respective keto form.

|  | 0*e* | 0*k* | 1 | 2 |
| --- | --- | --- | --- | --- |
| ES | -31.2 | -22.8 | -47.7 | -104.0 |
| POL | -90.0 | -83.1 | -43.3 | -19.5 |
| CT | -0.3 | -0.3 | -1.6 | -2.2 |
| REP | 91.8 | 90.1 | 89.7 | 90.0 |
| DISP | -42.5 | -42.7 | -45.7 | -47.1 |
| SOLV | 35.3 | 21.4 | 5.9 | 34.4 |
| INT | -37.0 | -37.3 | -42.7 | -48.4 |
